# Human goal-directed behavior is resistant to interventions on the action-outcome contingency

**DOI:** 10.1101/2025.04.26.650713

**Authors:** Omar David Perez, Sarah Oh, Anthony Dickinson, Jose Arenas, Emiliano Merlo

**Affiliations:** Department of Industrial Engineering, University of Chile, Santiago, Chile; Department of Psychology, University of California, Berkeley, California, USA; Division of the Humanities and Social Sciences, California Institute of Technology, _Pasadena_, California, USA; Instituto Sistemas Complejos de Ingeniería (ISCI), Santiago, Chile; Department of Psychology and Behavioural and Clinical Neuroscience Institute, University of Cambridge, Cambridge, UK; School of Psychology, University of Sussex, _Falmer_, UK

**Author notes:** Correspondence regarding this article should be sent to Omar David Perez or Emiliano Merlo. Authors contributed equally to this work (Omar David Perez, Sarah Oh).

**Keywords:** goal-directed, extinction, non-contingent learning, humans

## Abstract

Goal-directed actions are those performed with the expectation of producing a specific outcome and are therefore sensitive to changes in the outcome’s current value without re-experiencing the action-outcome contingency. Disrupting the action-outcome association by a number of interventions can reduce responding, but whether such interventions also affect the goal-directedness of human responses remains unknown. We trained participants to perform different actions for two distinct outcomes in two groups. One group experienced extinction (i.e., complete suspension of outcome delivery for a specific previously rewarded action) whereas the other was given non-contingent training (i.e., outcomes were presented independently of responding). Goal-directed control was tested by outcome devaluation, making one of the two outcomes less desirable, and responding was then tested in the absence of outcome presentations. Despite having been significantly reduced in both groups by extinction and non-contingent training, responding remained sensitive to outcome value, suggesting that human goal-directed behavior is resistant to action-outcome contingency interventions. We discuss these results in light of recent theories of goal-directed behavior.

## Introduction

Humans, like other animals, rely on multiple systems to guide their actions. One prominent distinction is between a goal-directed system, in which actions are selected based on their expected consequences, and a habitual system, in which behavior is driven largely by learned associations between cues and actions, without reference to the current value of the outcome. These systems are dissociable both computationally and neurally, and their interplay plays a role in many aspects of adaptive behavior, prompting several dual-system theories of learning and decision making (Daw et al., 2005; Daw & O’Doherty, 2013; Dolan & Dayan, 2013; Everitt & Robbins, 2005, 2016; Henriquez-Jara et al., 2025; Lee et al., 2014; Perez & Dickinson, 2020; Yin & Knowlton, 2006).

A critical feature of goal-directed control is its sensitivity to changes in outcome value. In classic outcome devaluation paradigms, individuals learn to associate specific actions with rewarding outcomes. The value of one outcome is then reduced by satiating the animals on that outcome or by pairing it with an aversive consequence. Behavior is then tested in the absence of outcome feedback (so that no new learning is possible). If an action is goal-directed, its selection will reflect the updated value of the outcome, being less likely to be selected if the outcome has been devalued. By contrast, responding will persist regardless of devaluation of the outcome when habits control behavior. This dual-system distinction has proven critical for understanding both healthy and impaired learning and decision-making, and has been used extensively in neuroscience and psychology to study compulsive and addictive behavior (Daw et al., 2005; Everitt & Robbins, 2005, 2016; Gillan et al., 2014; Lee et al., 2014; Tricomi et al., 2009; Yin & Knowlton, 2006).

Much of the evidence for these distinct forms of behavioral control comes from free-operant experiments in rodents, where subjects have the opportunity to perform an action freely, without any cues signaling when an action-outcome contingency is in effect. Seminal work by Adams (1982) showed how different amounts of training affected whether free-operant responding would be goal-directed or habitual. He trained hungry rats to press a lever in two different groups, a moderate training group that performed 100 responses under continuous reinforcement (all responses were rewarded) and another, extended training group that had 500 reinforced responses and therefore 5 times the training of the moderate training group. Adams (1982) found that habits only developed in the extended training group, while the group given moderate training remained goal-directed. Later on, Dickinson and colleagues established the response-outcome reinforcement schedule as critical in the development of habits. Ratio schedules, in which each response has a fixed probability of being reinforced, tend to be less likely to produce habits compared to interval schedules, where the probability is associated with the time elapsed between reinforcers (Dickinson, 1985; Dickinson et al., 1983). One prominent hypothesis for this difference is that animals track a correlation between the rate of responses and reinforcers, which is stronger in the ratio than in the interval case.

Most dual-system theories incorporating goal-directed and habitual systems assume that the strength of the action-outcome association should affect the degree to which outcome value affects responding. Indeed, since goal-directed behavior is sensitive to the expected value of outcomes—or in other words, to the likelihood of an action producing an outcome as well as the value of that outcome—any intervention on the action-outcome association should moderate the effect of outcome devaluation on responding. Specifically, responding should be less sensitive to outcome devaluation when action-outcome contingencies have been weakened. In folk psychology terms, a person who encodes that their actions are not strongly related to the likelihood of obtaining an outcome should be less sensitive to changes in that outcome value when deciding to act to obtain them (Daw et al., 2005; Perez & Dickinson, 2020; Perez & Urcelay, 2025).

Prof. Rescorla (1993) tested this prediction using the design laid out in Table 1 (Extinction condition). Initially, he trained hungry rats to perform four different instrumental responses with two of the responses (R1 and R3) being reinforced with one food outcome (O1), and the other two responses (R2 and R4) with a second food outcome (O2). Having established responding, two of the responses (R1 and R2) were extinguished by omitting their respective outcomes. At issue was whether this extinction had weakened the R–O association mediating goal-directed control. However, in order to assess such control with an outcome devaluation test, responding first had to be re-established using a third outcome (O3) such that the level of responding was comparable between extinguished and non-extinguished responses at the outset of the final test. In this test, Rescorla devalued O1 by conditioning a food aversion to it. Finally, the animals were given a choice between R1 and R2 and between R3 and R4 in the absence of any outcomes. To the extent that responding is under goal-directed control, the rates of R2 and R4 during the test should be higher than those of R1 and R3, respectively, given that the latter’s outcome, O1, is devalued. Moreover, if extinction reduces the strength of the R1-O1 and R3-O1 association, and hence goal-directed control, the difference between the R1 and R2 rates should be reduced relative to the difference between R3 and R4 rates.

**Table 1.**
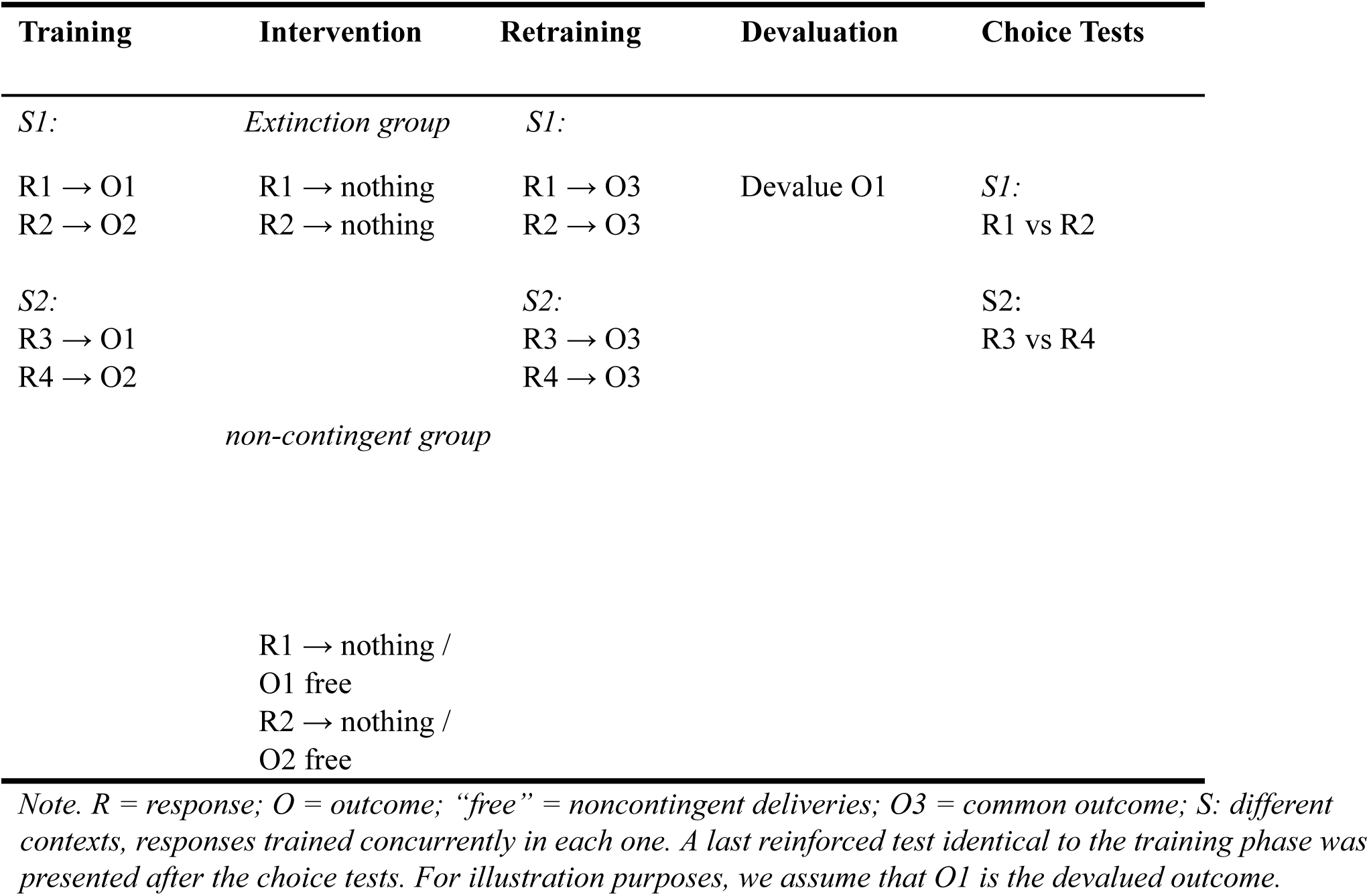
Procedure overview for extinction group / non-contingent group (based on Rescorla, 1993) R and O denote responses (keypresses) and outcomes (fictitious coins), respectively. S1 and S2 indicate different contexts, with different pairs of keys available to be pressed.

The fact that R3, originally trained with the now-devalued O1, was performed less on test than R4, trained with the still-valued O2, confirmed that the initial instrumental training established goal-directed control over responding. Surprisingly, however, the magnitude of the outcome devaluation effect revealed that the contrast between R1 and R2 was similar to that observed for R3 versus R4. Extinction of R1 and R2 by the total removal of their training outcomes had no effect on the extent to which the current value of these outcomes impacted performance, even when those actions no longer led to the rewarding outcomes and rats were not responding to obtain them by the end of the extinction phase.

Perez and Dickinson’s (2020) theory of free-operant behavior is unique in explaining why extinction does not impact subsequent goal-directed control. According to their theory, a mnemonic system tracks recent experience in short working-memory samples, computes the local correlation between action and outcome rates within each sample, and maintains a running average of this rate correlation. Goal-directed strength, which directly contributes to response rate, is taken to be g = I x r, the product of the correlation r and the current incentive value I of the outcome (Perez & Dickinson, 2020). Two predictions follow. First, during extinction, when no outcomes are present in incoming memory samples, the correlation can no longer be updated, so the training-established r is carried forward. Therefore, devaluation sensitivity should be preserved even as performance declines^1^.

A second prediction of rate-correlation theory is that degrading an action-outcome contingency by delivering non-contingent outcomes should decrease the experienced correlation r and, since g = I x r, the impact of revaluation (a change in I) on behavior should be attenuated. Recent work by Crimmins et al. (2022) in animals has investigated this prediction using a contingency degradation procedure: they delivered non-contingent outcomes at the same rate during periods when subjects did not perform a lever press, effectively establishing a zero correlation between lever pressing and outcome delivery (Dickinson & Mulatero, 1989; Hammond, 1980). Contrary to the prediction of rate correlation theory (Perez & Dickinson, 2020), they found that non-contingent training had no effect on the magnitude of outcome devaluation effect, at least when the instrumental training procedure was the same as that employed by Rescorla (1993) in that each response was trained on a separate manipulandum in separate sessions. However, when they employed a bidirectional manipulandum that controlled for Pavlovian confounds that may affect motivation and responding to a devalued outcome, they found that non-contingent training weakened the devaluation effect.

The question of primary interest in this study is whether the resistance of goal-directed control to extinction and contingency degradation observed in rodents extends to humans. To control for the Pavlovian confounds reported by Crimmins et al. (2022), we used a concurrent training procedure with matched stimuli, ensuring that Pavlovian associations were equally distributed across outcomes. This design tests Perez and Dickinson’s model and determines whether human goal-directed action is controlled by the experienced action-outcome rate correlation or by more complex causal representations.

## Methods

### Participants and apparatus

We recruited participants from the Prolific online platform (www.prolific.com). The task was programmed in JavaScript using the JsPsych library (De Leeuw et al., 2023). As there were no previous experiments testing the impact of outcome revaluation on human free-operant responding after degradation of the action-outcome contingency, we did not perform a power analysis before data collection. However, we aimed to reach 100 participants per group, which is considerably higher than similar investigations of human action learning (Perez, 2021; Reed, 2015). Participants with partial data were excluded from the final dataset. The final number of participants was 226, with 117 participants in the extinction group (mean age=41.07, sd=12.92, 46% females) and 109 participants in the non-contingent group (mean age=37.06, sd=12.15, 58% females). We excluded participants only if they stopped responding altogether in more than one training block. No participants met this criterion.

Participants were required to be located in the US or UK, fluent in English, and were paid a flat rate of 8 GBP per hour, plus a bonus payment ranging from 1-3 GBP depending on the number of coins obtained during the task (mean bonus group Extinction: 2.81 GBP; group non-contingent: 2.65 GBP). The study was approved by the University of Sussex Sciences & Technology Cross-Schools Research Ethics Committee (SCITEC) (approval ER/EM540/1).

### Procedure

Our design was an adaptation of the general paradigm originally developed by Rescorla ^11^. The procedure implemented the general design outlined in Table 1 with the instrumental contingency being degraded by omitting all outcomes in the extinction group, and by presenting outcomes independent of responding in the non-contingent group. At the beginning of each experiment, participants were told that they should press keys to obtain coin outcomes of two different types, red and blue (O1 and O2), and that their goal was to obtain as many coins as they could. They were also told that there was a third type of coin, green, which could also be earned in some blocks of the experiment, and that this coin was worth the same as the other two (O3 in Table 1). Participants were also informed that the collected coins would be deposited in different fictitious piggy banks of limited size, one for each type of coin.

Two actions (R1 and R2) were exposed to the contingency intervention (extinction or non-contingent training), whereas the other two actions (R3 and R4) did not undergo any intervention and functioned as within-subject controls. In each training block, participants had the opportunity to press two keys, either the two top keys, “q” or “p”, or the two bottom keys, “z” or “m”. The location of the keys on the screen respected their relative position on the keyboard (e.g., “z” was shown on the bottom left side of the screen whereas “p” was shown on the top right location). For each participant, one of the top keys delivered red coins and the other one blue coins. To avoid participants associating the type of coin with the side of the response (which would have associated the two outcomes to different hands) the opposite assignment was programmed for the bottom keys. Therefore, actions performed with one of the hands always led to different outcomes across different blocks of training. For example, if for one participant the assignment for top keys was q *→* red and p *→* blue, then for the next block the assignment was z *→* blue and m *→* red.

The sequence of stages in the procedure is illustrated in Figure 1. Initially, participants underwent three 120-s blocks of concurrent training for each pair (“q” and “p”, “z” and “m”) of actions. The starting pair of actions was randomized across participants and the blocks were presented sequentially once this first assignment was set for each participant. Unlike the original design of Rescorla, where subjects were trained in different sessions each with a different response, in our design two actions were concurrently trained under a random-interval (RI) schedule (Kosaki & Dickinson, 2010), so that on average the appropriate coin for each response became available to be collected every T seconds. This was implemented by programming two independent clocks, each setting up an available outcome with a probability of 1/T per second. To minimize accidental reinforcement of a heuristic strategy that alternates between responses, we introduced a changeover count: only the second response after a switch could produce a coin, even if the schedule on the new action had already an available outcome while the participant was engaged in the other response. The parameter T was set to 15s for both the extinction and non-contingent groups.

**Figure 1.**
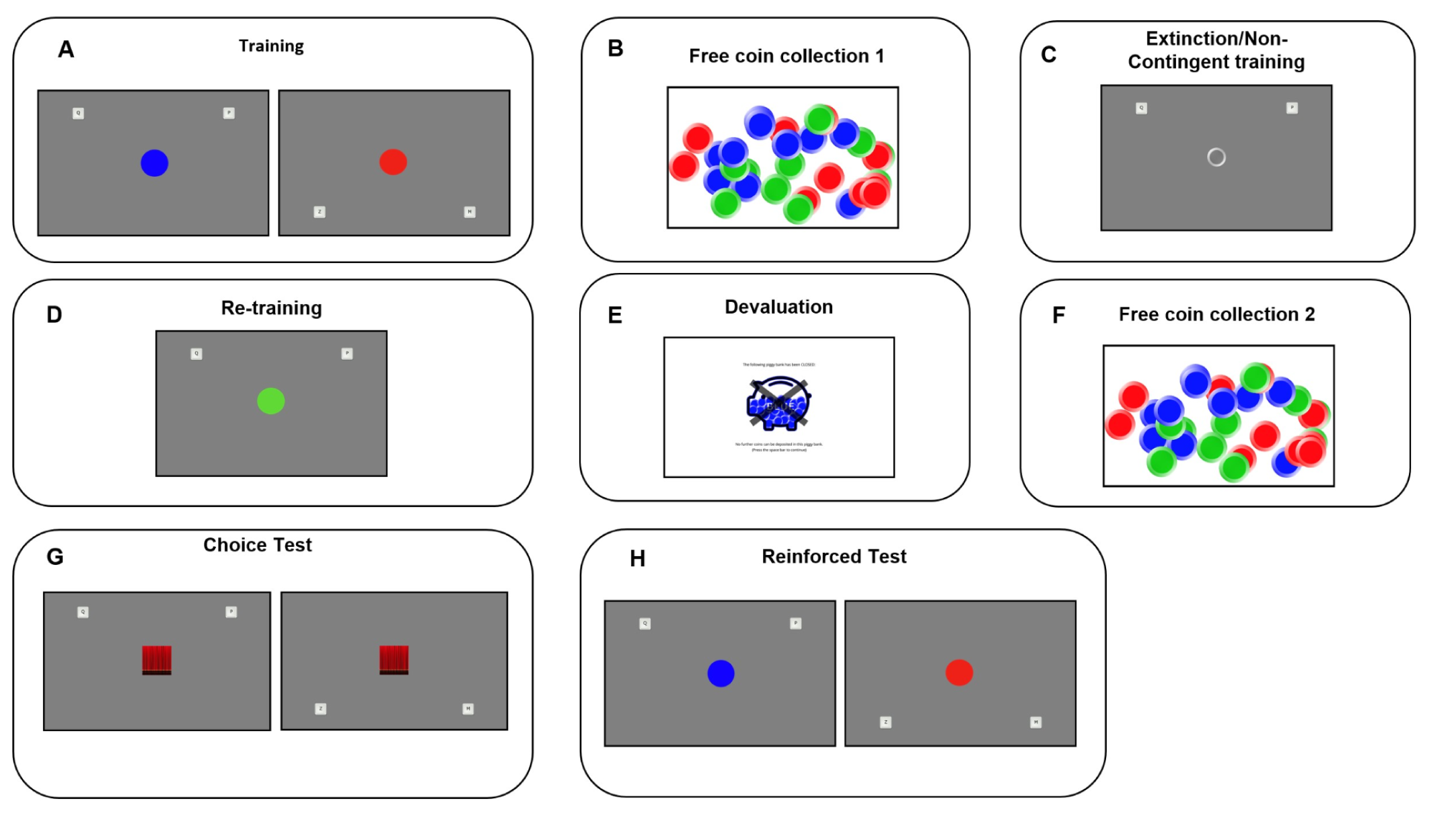
Task timeline, screens, and design. Participants learned to press four keys to earn two token outcomes (red or blue coins). Keys were trained in concurrent pairs under RI-60s schedules—the top keys “q”/“p” and the bottom keys “z”/“m”—and the outcome–key mapping was reversed across top vs. bottom pairs for each participant to avoid confounding outcome identity with hand/side. Training used independent, identical random-interval (RI) schedules per key. **(A) Training.** Two contexts with two keys each; each key produced a distinct token (red/blue). **(B) Free coin collection 1.** Baseline consumption check. **(C) Contingency intervention.** For two actions (within-subject), the contingency was altered depending on group: Extinction (trained outcomes omitted) or Non-contingent (matched-rate outcomes delivered independently of responding). The other two actions served as controls. **(D) Re-training.** Responding re-established for all actions using a common “green” coin to equate rates prior to test. **(E) Devaluation.** One outcome (O1) was devalued (piggy bank “closed”). **(F) Free coin collection 2.** Post-devaluation check. **(G) Choice (“curtain”) test.** Participants chose between the two actions in each context with outcomes hidden but still earned, providing a feedback-free test of value sensitivity. **(H) Reinforced test.** Outcomes were visible and contingent exactly as during training, enabling comparison of value sensitivity with vs. without feedback. **Durations.** Training 1: 6×2 min; Training 2: 6×2 min; Contingency intervention: up to 15 min; Re-training: 6×2 min; Choice test: 4×1.5 min; Reinforced test: 2×2 min.

Once a total of three blocks of training for each pair of actions finished, participants were presented with a message informing them that the piggy banks were filling up (see “Devaluation” screenshot in Figure 1). This information was added to remind participants that their coins were being deposited and counted during the experiment and to familiarize them with the stimuli (indicating a full piggybank) that would be presented at the end of training after devaluation. After this screen, participants were presented with 30 coins located in random positions on the screen, 10 of each type, and asked to collect a total of 10 coins that would be deposited into their respective piggy banks.

The free consumption phase (“Free coin collection” in Figure 1) was followed by a 15-min extinction or non-contingent stage wherein two actions (either “q” and “p”, or else “z” and “m”) were presented and outcomes were omitted in the extinction group or followed by the non-contingent presentation of both outcomes each on an independent random interval 15-s schedule in the non-contingent group. On this latter schedule, participants could achieve the same outcome rate irrespective of whether they responded or not, thereby generating a zero contingency between action and outcome^2^.

After the intervention phase, participants were given the opportunity to perform a 120-s block of retraining of all actions, which was similar to those in the training phase except for the fact that the outcome was a green coin for all responses. Participants were informed that these coins were worth the same as the blue and red ones, and that they would also be deposited into their respective piggy banks. In this manner, we ensured that participants were performing responses at comparable rates at the end of training despite the fact that extinction or non-contingent training should have reduced responding during the intervention phase. This retraining phase was followed by devaluation of one of the outcomes. In this phase, participants were presented with a screen showing one of the piggy banks being filled up so that no more coins of that type could be deposited in the piggy bank, effectively devaluing that outcome with respect to the others. To test if the devaluation phase succeeded in achieving this goal, we then presented participants with another consumption test where they were asked to collect 10 coins using mouse clicks in the same manner as the previous coin collection phase (see Figure 1). During this free collection phase, we expected that participants would collect less of the devalued coin than the two other still-valued coins. This would suggest encoding of outcome value and effectiveness of the devaluation procedure in reducing it.

Participants were then given the critical outcome devaluation test, at the start of which they were told that they could respond to obtain coins in the same manner as in the training phase for 90 s, but that this time the outcomes obtained would be covered by a curtain (pseudo-extinction choice test). They were informed that, apart from that, nothing about the experiment had changed. The coins obtained would be still deposited into their respective piggy banks (provided they were not full). Finally, participants underwent an additional reinforced test in which the curtain was removed and the outcomes were again presented contingent on the actions as in the training phase.

Once the behavioral task finished, to test their explicit knowledge of the action-outcome contingencies, the participants were presented with the outcomes on the screen and asked to press the actions that led to each of the two types of coins. The number of correct answers ranged from 0 (no correct action-outcome mapping identified correctly) to 4 (all action-outcome mappings identified correctly).

## Results

All our analyses were performed in the R programming language under the RStudio IDE (RStudio Team, 2020). We fit a mixed-effects model with fixed effects of Group via *lmer* (extinction vs non-contingent), Value (valued vs devalued), Contingency (degraded/extinguished vs control), and all interactions, with random intercepts and by-subject random slopes for value. The primary dependent variable was response rate during the test phases, expressed as responses per minute (computed from n_resp_stage and the fixed phase duration). Analyses used the following factors and reference levels: Group (between-subjects: Extinction vs Non-contingent; reference = Extinction), Contingency (within-subjects: control vs intervened; reference = Control), Value (within-subjects: valued vs devalued; reference = valued), and, where tests were compared, Stage (within-subjects: Choice test vs Reinforced test; reference = Choice test). The devaluation effect was operationalized as the within-subject contrast valued − devalued estimated from mixed-effects models with random intercepts and by-subject random slopes for Value.

Our primary comparison tested whether the devaluation effect differed by Group in each test phase by contrasting a full model (response ∼ Value * Contingency * Group + (1 + Value | ID)) with a reduced model (response ∼ Value * Contingency + Group + (1 + Value | ID)). This contrast was evaluated with a likelihood-ratio test and also summarized with a BIC-approximate Bayes factor. For other reported effects, estimates come from the corresponding mixed models; where nested comparisons were required, we used likelihood-ratio tests. We also evaluated a more complex “maximal” model that included random slopes for Contingency, but this model resulted in a singular fit, indicating that the data did not support the additional complexity (likely due to low variance in the individual responses to the contingency manipulation).

To characterize within-phase changes in responding, we fit a separate mixed-effects model on the intervention-phase data binned into successive 3-min intervals. The dependent variable was the total number of responses per bin. Fixed effects included Group (Extinction vs. Non-contingent), Bin (coded 1-5 and mean-centered to capture a linear trend), and their interaction. Given the relatively small number of observations per participant (five bins), we specified random intercepts only. This parsimonious random-effect structure was chosen to avoid over-parameterization and ensure model convergence, as the primary purpose of this specific analysis was to summarize and compare the general rate of decline across groups rather than to estimate individual differences in extinction rates.

### Training

Figure 2A shows the final mean response rates during the last 2 min of each phase of the extinction and non-contingent groups. A Welch’s t-test was used to confirm comparable performance across groups before contingency degradation. No group difference was detected in the last 2 min of the last training block, with extinction (M = 119.28 responses per min) and non-contingent (M = 109.61 responses per min) performing similarly (t(221.95) = 1.24, p = .218, mean difference = 9.67, 95% CI [−5.76, 25.11]), confirming comparable performance prior to the contingency intervention phase. In fixed-duration RI blocks, participants produced on the order of 200–250 actions and 10–13 outcomes per 2-min training/retraining block, and ∼45–74 rewards over the 15-min intervention, confirming ample exposure to the programmed contingencies to support learning (Tables 2 and 3).

**Figure 2.**
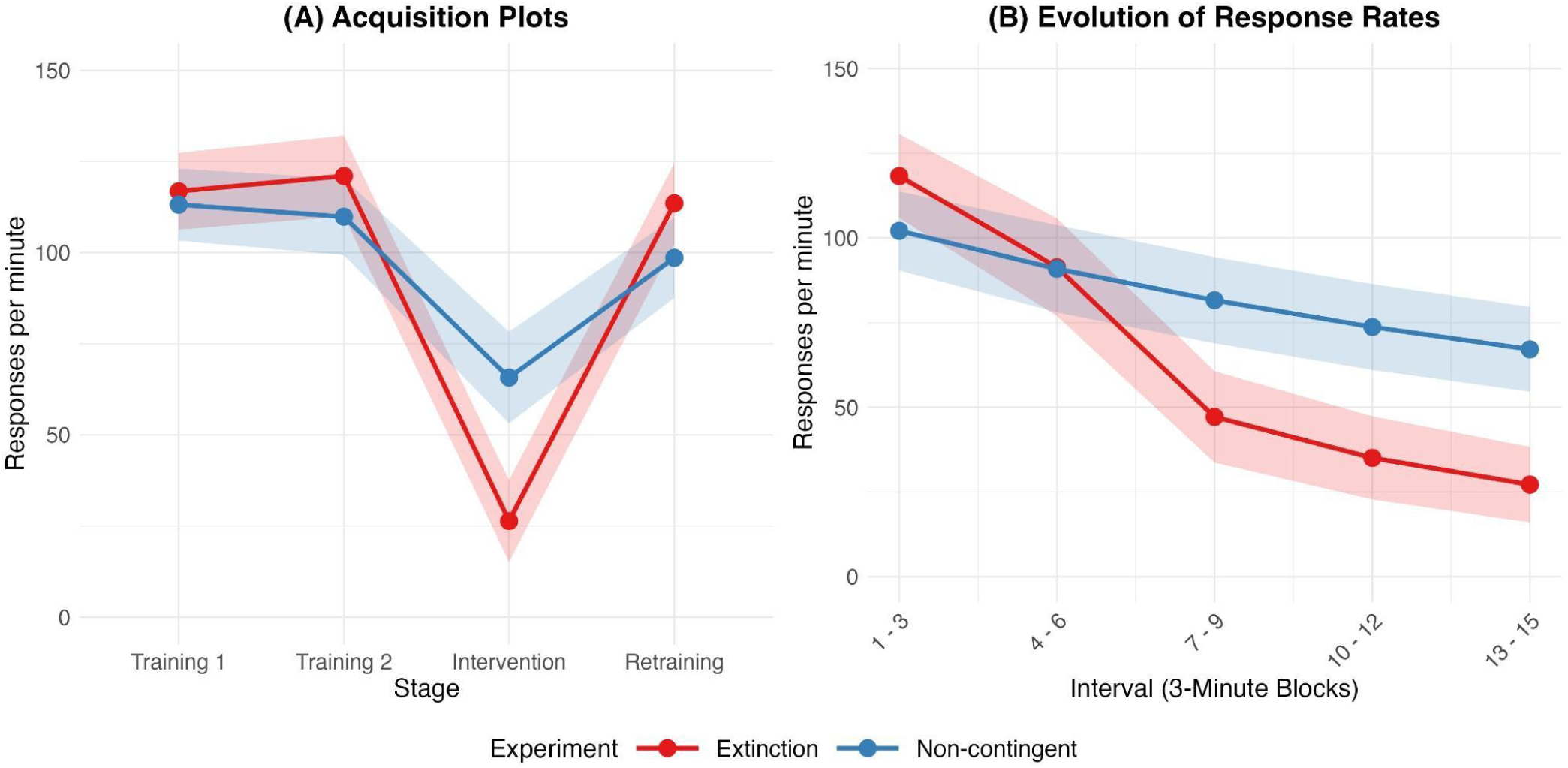
Response rates for training and during the intervention phase. **A)** Mean response rates during training, intervention and retraining phases for the extinction and non-contingent groups. Means are calculated for the last 2 minutes of each phase. **B)** Mean response rates during the intervention phases. Means are calculated for each of the five 3-min blocks. Error bands are standard errors of the mean.

**Table 2.** Average Number of Responses per Phase by Group.

| Group | Training<br>1 | Training<br>2 | Choice<br>Test | Intervention | Retraining | Reinforced<br>Test |
| --- | --- | --- | --- | --- | --- | --- |
| Non-contingent | 233.18<br>(8.75) | 223.52<br>(10.24) | 162.59<br>(9.35) | 1267.77<br>(87.67) | 195.65<br>(10.72) | 195.53<br>(11.13) |
| Extinction | 238.82<br>(8.92) | 248.09<br>(10.36) | 197.83<br>(8.88) | 984.05<br>(81.84) | 229.21<br>(10.32) | 228.52<br>(10.62) |

Note. Entries are $M (SE)$ .

**Table 3.**
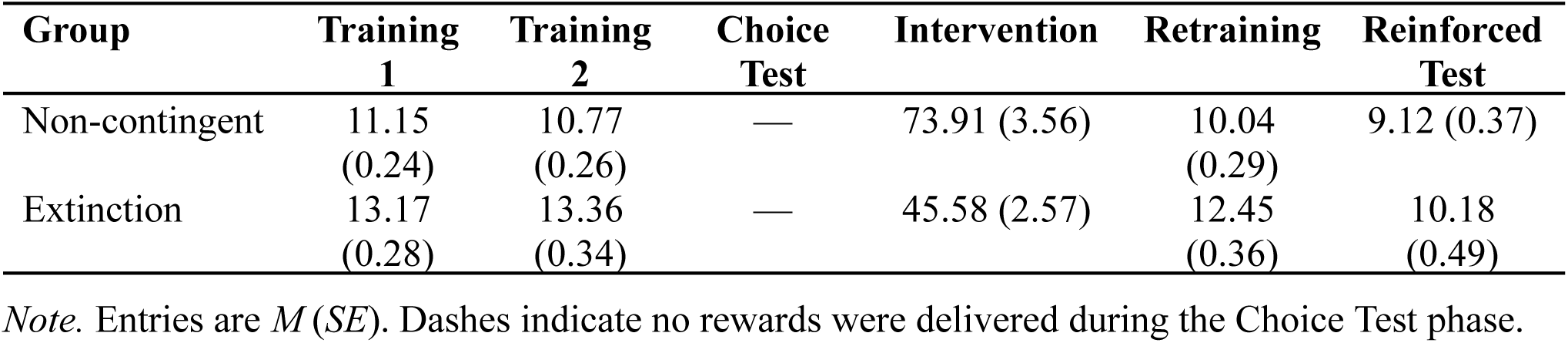
Average Number of Rewards per Phase by Group.

| Group | Training<br>1 | Training<br>2 | Choice<br>Test | Intervention | Retraining | Reinforced<br>Test |
| --- | --- | --- | --- | --- | --- | --- |
| Non-contingent | 11.15<br>(0.24) | 10.77<br>(0.26) | — | 73.91 (3.56) | 10.04<br>(0.29) | 9.12 (0.37) |
| Extinction | 13.17<br>(0.28) | 13.36<br>(0.34) | — | 45.58 (2.57) | 12.45<br>(0.36) | 10.18<br>(0.49) |

Note. Entries are $M (SE)$ . Dashes indicate no rewards were delivered during the Choice Test phase.

The evolution of response rates across the intervention phase exhibited a clear decline in both groups over successive 3-min bins (see Figure 2B; main effect of Bin: b = –73.6, SE = 3.33, p < .001). However, extinction produced a sharper reduction than non-contingent training. Although the non-contingent group started at lower overall levels (Group effect: b = –84.2, SE = 27.8, p = .003), their decline over time was weaker, as revealed by a significant group x bin interaction (Group x Bin: b = 47.0, SE = 4.61, p < .001). Thus, by the end of the intervention phase, participants in the non-contingent group maintained relatively higher response rates compared to those in the extinction group, consistent with the slower suppression typically reported for non-contingent procedures (Rescorla & Skucy, 1969).

### Choice test

Figure 3A shows the mean response rates during the choice test for each group. In both groups, participants responded less for actions associated with the devalued outcome than for those associated with a still-valued outcome. The analysis revealed a significant main effect of Value (b = −19.2, SE = 4.59, p < .001), indicating a robust reduction in responding for the devalued outcome. However, there was no main effect of Contingency (b = 1.17, SE = 1.96, p = .549), and no significant interaction between Value and Contingency (b = 2.16, SE = 3.92, p = .581). The interaction between Value and Group was also not significant (b = 14.8, SE = 9.19, p = .109), nor was the three-way interaction among Value, Contingency, and Group (b = −7.06, SE = 7.83, p = .368). These results indicate that the magnitude of the outcome devaluation effect did not reliably differ between Extinction and Non-contingent training groups. A likelihood-ratio test comparing the full model to a reduced model without the three-way interaction showed no improvement in fit (chi-sq(3) = 3.43, p = .330). A BIC-approximate Bayes factor provided extreme evidence in favor of the reduced model (BF01 = 4,890), confirming that the devaluation effect did not differ across intervened actions or groups.

**Figure 3.**
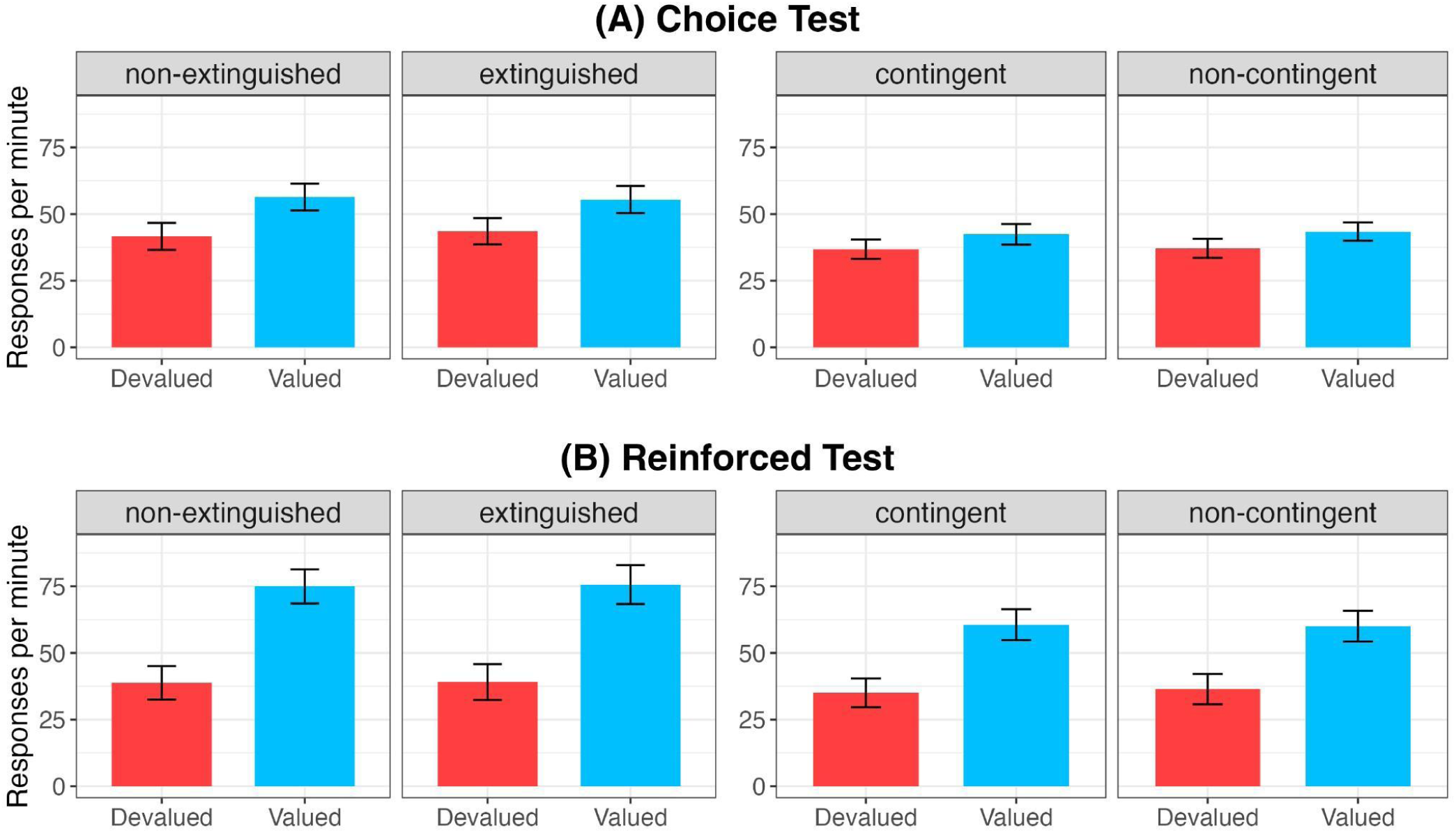
Response rates in the tests for extinction and non-contingent groups. **A)** Average responses rates during the choice tests. **B)** Average response rates during the reinforced tests. Error bars represent 95% within-subject confidence intervals (Morey, 2008).

### Reinforced test

Figure 3B shows the response rates during the reinforced test, in which the outcomes were reintroduced contingent on responding, identical to the training phase. The same value-based pattern was observed, with a larger devaluation effect overall (Value: b = −61.0, SE = 6.93, p < .001). As in the choice test, neither Contingency nor its interaction with Value was reliable (b = 0.899, SE = 2.23, p = .687; Value x Contingency: b = 1.58, SE = 4.46, p = .724), and the three-way interaction with Group was not significant (b = 4.62, SE = 8.92, p = .604). The reduced model was again strongly preferred (χ²(3) = 3.14, p = .370), with a Bayes factor providing extreme evidence in favor of the more parsimonious model (BF_01_ = 5,640). Critically, a combined analysis across the two tests revealed that the devaluation effect was significantly stronger in the reinforced test than in the choice test (Value x Phase: b = 46.0, SE = 5.33, p < .001), and this enhancement of the devaluation effect did not differ by Group (Value x Phase x Group: b = 8.68, SE = 7.41, p = .242; χ²(3) = 4.88, p = .181). The Bayes factor again provided extreme evidence against a group difference in this enhancement (BF_01_ = 6,720).

### Behavioral suppression and contingency knowledge as moderators

Because individual differences in attentional engagement/strategic adaptation during the intervention and in explicit contingency knowledge could each shape value sensitivity, we added two subject-level predictors to the test models: suppression (last 2 min of Training 2 minus last 2 min of the intervention; higher = more suppression) and contingency knowledge (contingency score; the number of action-outcome mappings correctly identified at the end of the task by each subject, ranging from 0 to 4). Each of these variables interacted only with value, while keeping the original group x contingency x value variables in the model. These two additional predictors were statistically independent (r = −.01, 95% CI [−.145, .116], p = .830). In the choice test, both independently moderated the value effect (Devalued vs Valued): Value x Suppression (b = −0.133, SE = 0.06, p = .033; Value x Contingency Score b = −21.2, SE = 3.46, p < .001). The same pattern held in the reinforced test, with larger coefficients (Value x Suppression b = −0.270, SE = 0.09, p = .003; Value x Contingency Score b = −37.6, SE = 5.01, p < .001). In all cases greater suppression and higher contingency knowledge were associated with a stronger devaluation effect, and these relationships held over and above the other effects we report.

## Discussion

Our experiment examined whether degradation of the action-outcome contingency reduced the goal-directed control of human action as assessed by the outcome-devaluation test. In the Extinction group, both extinguished and non-extinguished actions were value-sensitive, which is consistent with the original results obtained by Rescorla (1993) in rodents. Similarly, the Non-Contingent group replicated in human participants the Crimmins et al. (2022)’s finding in rodents that the degradation of the instrumental contingency by training with non-contingent outcomes can also have no impact on the outcome devaluation effect, at least when the actions involve responses on different manipulanda (levers in the case of rodents and key presses in the present human experiment).

The theoretical interpretation of these findings is complex. The theory offered by Perez and Dickinson (2020) is unique in providing an account of why goal-directed control is resistant to extinction. According to this theory, it is the mean local experienced correlation between response and outcome rates in working memory that determines the strength of goal-directed control. This computational memory system predicts that extinction should have only a limited impact on goal-directed control. Because the rate correlation can only be computed when both responses and outcomes are represented in the working memory and, by necessity, the computation is rendered impossible when the memory is cleared of representations of at least one of the event types (either the response or the outcome) this suspension should happen relatively early in extinction with the absence of any outcome representation. Thereafter the strength of goal-directed control (g) will reflect the mean running average rate correlation (r) established earlier during training, with no further updates. Since g = I x r, and r is still positive, changes in the incentive value of the outcome (I) should be immediately manifest in behavior. Perez and Dickinson (2020) verified this incidental prediction of the theoretical model by formal simulation.

A challenge to Perez and Dickinson’s rate-correlation theory is that non-contingent training failed to reduce goal-directed control in our study, a finding that mirrors recent rodent data (Crimmins et al., 2022). Perez and Dickinson explicitly predicted that non-contingent outcomes should degrade goal-directed control more effectively than extinction; however, our results and others suggest otherwise.

One possible explanation for this persistence is Pavlovian mediation. Classic two-process theory (Rescorla & Solomon, 1967) suggests that stimuli (like levers or buttons) can form Pavlovian associations with outcomes, allowing them to drive responding independently of the action-outcome contingency. Crimmins et al. (2022) demonstrated that when a manipulandum predicts a specific outcome (e.g., a single lever in a specific session), Pavlovian influences can mask the degradation of goal-directed control. When they used a bidirectional lever that equally predicted both outcomes—thereby neutralizing Pavlovian bias—their results aligned with rate-correlation theory.

To control for these confounds in our online setting, we utilized concurrent training so that the stimuli accompanying each pair of responses were equally associated with both outcomes. While we acknowledge that the use of different buttons may still be susceptible to subtle Pavlovian influences, the robust value-sensitive responding observed here represents a significant challenge to the memory-based tracking of rate correlations. This resilience suggests that human goal-directed control may be mediated by explicit causal representations that are less sensitive to contingency fluctuations than traditional models assume (Perez & Soto, 2020).

These findings suggest a need to refine the theory to account for human-specific cognitive strategies. While humans show sensitivity to rate correlations in their underlying response rates, their explicit causal reasoning is more closely tied to the probability of reinforcement. As Perez and Soto (2020) found, instrumental performance can be higher under schedules with high rate correlations (such as ratio schedules), yet causal beliefs can correlate with reward probability per action. This indicates that humans may rely on explicit causal reasoning beyond memory-based rate tracking. This discrepancy provides a constructive opportunity to refine rate-correlation models by incorporating explicit causal variables, acknowledging that goal-directed action in humans is mediated by a causal representation that is distinct from the processes governing response frequency.

Although both contingency interventions significantly reduced responding, it is notable that extinction produced a greater reduction than non-contingent training (Rescorla & Skucy, 1969). It is also notable that there was still considerable responding after both contingency interventions. However, we think it is unlikely that greater reductions in responding would have decreased the outcome devaluation effects. Further analyses incorporating the magnitude of response reduction during extinction and non-contingent training revealed a significant suppression x value interaction, such that participants who showed greater reductions in responding also exhibited stronger outcome devaluation effects. Importantly, this effect was independent of participants’ explicit contingency knowledge, which also predicted value sensitivity but was not correlated with suppression, suggesting that the two variables reflect distinct psychological processes contributing independently to goal-directed control. These results probably reflect individual differences in the cognitive resources applied to the task—the greater the resource allocation to encoding the relative contingencies, the more likely it is that a participant will have reduced responding under extinction and non-contingent training and learned the response–outcome associations and outcome values necessary for the devaluation effect. This suggests that while suppression may capture attentional engagement or strategic adaptation during the intervention phase, contingency scores appear to reflect explicit knowledge of instrumental structure. And they both appear to contribute independently to goal-directed control.

We observed a stronger devaluation effect in the reinforced test compared to the choice test, a pattern that likely reflects differences in how action-outcome representations are accessed and expressed. According to rate-correlation theory, goal-directed strength depends on the perceived correlation between actions and outcomes. In the choice test, the absence of outcomes (covered by a curtain) may effectively “freeze” this representation at its last experienced training value, limiting the behavioral expression of the devaluation. In contrast, the delivery of outcomes during the reinforced test allows for the active reactivation or ongoing computation of this correlation in working memory, which can amplify the expression of goal-directed control. Furthermore, receiving outcomes provides direct experiential feedback that likely enhances the incentive salience and discriminability of the outcome’s current value. This feedback may also increase overall motivational engagement or attention to the specific contingencies, facilitating a more strategic expression of goal-directed behavior compared to the purely memory-based Choice Test.

Our paradigm is the first free-operant experiment in humans including a reinforced test at the end of training. Previous studies in humans have ignored this fundamental test in assessing the effect of value on responding after interventions aimed to prompt habit formation and relied either on consumption tests similar to our coin collection procedure or specific satiety when primary reinforcers are involved (Gillan et al., 2015; Pool et al., 2019, 2022). Although such a test is necessary to show the effectiveness of devaluation, it is not sufficient: what is more critical is showing that in addition to that sensitivity, revaluation of an outcome is effective in the same training context and contingent on the actions performed during training. Our data show that the devaluation method was effective in doing so by revealing a stronger effect of outcome value in the reinforced choice test than in the pseudo-extinction choice test.

This study is not without limitations. First, while participants’ behavior indicates that outcome value was encoded—as shown by the stronger effect of value on the reinforced test than in the choice test—the use of artificial tokens may not provide as strong an incentive as primary reinforcers or direct monetary payments, potentially limiting motivation. Second, although longer interventions are unlikely to change the main devaluation results, the fact that non-contingent training reduced responding less strongly than extinction may have limited the sensitivity to detect small interactions. Moreover, the influence of Pavlovian learning is difficult to discard in an online task, as the use of bidirectional manipulanda is difficult to implement without specialized hardware such as joysticks. Finally, while the online setting contributes to variability in engagement, it is worth noting that the inclusion of attention checks may disrupt the formation or expression of habits (Bouton, 2021, 2024; Thrailkill et al., 2018).

In addition to environmental factors, broader cognitive mechanisms likely contribute to the individual differences observed in goal-directed control. Indeed, variability in working memory capacity and general intelligence has been associated with the capacity to shift flexibly between habitual and goal-directed systems. Individuals with greater cognitive resources may be better equipped to maintain and update representations of action-outcome associations, thereby sustaining value-based control even when contingencies are degraded.

In spite of these limitations, the results point to a straightforward conclusion: weakening the action-outcome contingency through extinction-like procedures does not eliminate value-based control. Consider the implications for drug-seeking behavior. Making the drug less reliably available following an action—or arranging periods in which the action no longer produces the drug—is unlikely to reduce sensitivity to the drug’s value when it is encountered again. If the subject still values the drug, goal-directed choice will recover.

Interventions targeting outcome value should therefore be prioritized over those that merely degrade the action-outcome contingency. This aligns with approaches that modify incentive structure: contingency management that increases the relative value of abstinence, pharmacotherapies that reduce the drug’s hedonic or expected value, or policies that introduce immediate costs. By contrast, strategies that rely solely on weakening the action-drug association are unlikely to succeed.

In conclusion, our findings show that human goal-directed behavior is resistant to disruptions in action–outcome contingency and point to the robustness of goal-directed systems in humans. While extinction and non-contingent training reduce response rates, they do not eliminate value-based control. Our results extend those observed in humans for interventions of training extension, where goal-directed control has also been robust to different amounts of training (de Wit et al., 2018; Garr et al., 2021; Gera et al., 2023; Pool et al., 2022) and represent a challenge for Perez and Dickinson’s rate-correlation theory, which predicts different devaluation sensitivities for extinction and non-contingent training. It seems that, under many circumstances, human expectations of producing a valued consequence are remarkably difficult to erase.

## Declarations

## Abbreviations

BF: Bayes factor;
BIC: Bayesian information criterion;
CI: confidence interval;
IRB: Institutional Review Board;
RI: random-interval (schedule);
S: stimulus (or context);
R: response;
O: outcome;
SE: standard error.

## Ethics approval and consent to participate

This study was reviewed and approved by the University of Sussex Sciences & Technology Cross-Schools Research Ethics Committee (SCITEC), approval number ER/EM540/1. All participants provided informed consent prior to participation. All procedures were carried out in accordance with the ethical standards of the institutional and/or national research committee and with the 1964 Helsinki Declaration and its later amendments or comparable ethical standards.

## Consent for publication

Not applicable.

## Availability of data and materials

The dataset and analysis code that support the findings of this study will be deposited in a public repository (e.g., OSF/Zenodo). Access links will be provided upon acceptance and will be included in the published article.

## Authors’ contributions

ODP, SO, AD, and EM designed the study. SO programmed the tasks. ODP and JA analyzed the data. ODP and AD drafted the manuscript, with critical revisions from all authors. EM acquired funding. All authors read and approved the final manuscript.

## Acknowledgements

Not applicable

## Footnotes

1 The reader may wonder why responding is reduced by extinction if goal-directed strength is maintained. However, Perez and Dickinson (2020) embedded the goal-directed system within a dual-system theory that includes a habitual system. Within this dual-system account, extinction is primarily due to the development of response inhibition by the habitual system, which accords with the explanation offered by Rescorla (1993).

2 The use of a random time schedule to generate non-contingent outcomes differs from that employed by Crimmins et al (2022). They trained their rats on a ratio rather than interval schedule and therefore rendered the outcomes non-contingent by arranging that the outcome probability in a given 1s period in the absence of a response was the same as that following a response.

